# Cytocapsular tube network-tumor system is an integrated physical target for the highly effective and efficient pharmacotherapy of solid cancers

**DOI:** 10.1101/2022.01.19.476991

**Authors:** Tingfang Yi, Gerhard Wagner

## Abstract

Cancer is a leading cause of human lethality and cancer drug pan-resistant tumor metastasis (cdp-rtm) is a major source of cancer death. The integrated cytocapsular tube (CCT) network-tumor system (CNTS) is essential for cancer evolution procedures including cancer cell proliferation, migration and dissemination in CCT networks, new tumor growth in CCT terminals, conventional cancer drug pan-resistance, and tumor relapse. The preclinical screening experimentations with CCTs and networks are necessary prerequisites for the discovery and development of cancer drug candidates for efficient clinical trials, and effective and precise clinical cancer pharmacotherapy. However, it is unknown whether the popularly employed conventional mouse experimentations for preclinical therapeutic development generate CCTs and networks. Here, we comprehensively investigated the cancer cell line derived xenografts (CDX) of 8 kinds of cancer cell lines, and patient cancer cells derived-xenografts (PDX) with 16 kinds of clinical primary and metastatic malignant tumor cells in multiple cancer stages by immunohistochemistry staining assays with CCT marker protein antibodies of anti-CM-01 antibodies. We found that there are no CCTs or CCT networks in all these examined 16 CDX and 22 PDX transplanted tumors. Our data evidenced that the conventional transplantation tumors for preclinical therapeutic development do not engender CCTs or CNTS, which is consistent with and provides an explanation of the poor clinical outcomes of the marketed cancer drugs. This study demonstrated that CCT network-tumor system (CNTS) is an integrated physical target for pharmacotherapy, and that preclinical experimentations engendering CNTS (such as CCT xenograft, CCTX) should be employed for the discovery and development of highly effective and efficient cancer drugs aimed for cure of solid cancers.

## Introduction

Cancer is a leading source of human lethality worldwide. Conventional cancer drug pan-resistant tumor metastasis (cdp-rtm) is a major source of cancer death^1^. It is estimated that approximately 10 million cancer deaths and approximately 20 million new cancer cases occur in 2020 alone worldwide^2^. There are approximately 560 kinds of conventional cancer diagnosis methods but none can provide precise and decisive services for early-stage prognosis and diagnosis. Furthermore, US FDA approved about 650 cancer drugs since1949, and thousands of cancer remedies are employed around the world every day. The global cost of cancer was approximately $2.5 trillion in 2010 alone^3^. However, the marginal treatment outcomes suggest that the mechanisms of the majority of the nature of cancers are still beyond our understandings^3–6^.

Current hypotheses for cancer mechanisms, that seem plausible are actually controversial to deep clinical observations, will lead to inclined researches, indecisive diagnosis, misdiagnosis, development of poorly-efficient drugs, ineffective pharmacotherapy, huge cost, and poor outcomes. In the past 5,000 years, human have fought an intensive war against cancers. Multiple hypotheses on mechanisms of cancer metastasis were documented, including: seed-soil description^4^, metastatic cascade (with five steps of invasion, intravasation, circulation, extravasation and colonization), cancer stem cell (CSC), and so on^5–7^. In 1889, Paget described the distribution of secondary growth in breast cancer in botanical terms as the interaction of “seeds” (tumor cells) and “congenial soil” (the metastatic microenvironments)^4^. This metaphor from 130 years ago, like a “trunk” of a cancer hypothesis “tree”, largely influenced or derived multiple cancer metastasis hypotheses (“branches”) afterward. These hypotheses include metastatic cascade, cancer stem cells (CSCs), dormant metastasis, genomic instability driven cancer metastasis, “seeding” microenvironments in metastasis (mixed with extracellular matrix (ECM), immune cells, stromal cells and angiogenesis, matching “congenial soil”), and so on^8–19^. In all these hypotheses, metastatic cancer individual cells must overcome numerous physical ECM and neighbor cell obstacles barring metastasis and thus must invade vasculature to reach far distance destinations. However, all these hypotheses are controversial to clinical observations: 1) there are no circulating tumor cells in the blood in solid cancer stages I and II in which millions or hundreds of millions of cancer cells have already metastasized to far distance sites diagnosed as revealed with novel precise cancer diagnosis methods^20–22^; 2) all current commercial cancer drugs developed based on these hypotheses targeting solid cancers and cancer metastasis unexceptionally lead to conventional cdp-rtm and subsequential cancer death^1, 21–22^. It is necessary to know whether the conventional experimentations designed and based on these traditional hypotheses for preclinical therapeutic development generate CCTs and CCT networks.

Recently, we discovered that single cancer cells can engender superlarge, superlong, membrane-enclosed tube-shaped cytocapsular tubes (CCTs) outside of the cytoplasm membrane of cancer cells^20–22^. These previously invisible and unknown CCTs interconnect and from 3D superlarge networks across tissues and organs in cancer patients. CCTs and networks are propertied with pleiotropic biological functions: 1) provide physical freeway systems for cancer cell dissemination free of obstacles from ECM and neighbor cells, and for highly efficient cancer cell dissemination across tissue and organs, 2) CCT network systems are independent of the circulation systems (blood and lymph vessel systems) and free of the attack from immune cells and cancer drugs in the circulation systems; 3) CCT membranes function to protect cancer cells, to shield conventional cancer drugs outside, and reduce access to cancer cells in the CCT lumens; 4) damage-resilient CCT networks provide freeway systems and superdefences for the surviving cancer cells in CCT lumens; 5) the enlarged cytocapsulae function as stable ends for cancer cell proliferation, new tumor growth and formation, tumor relapse, and conventional cancer drug pan-resistance; 6) CCT freeway networks interconnect most/all tumors in the cancer patient’s body and form a potent, dynamic and integrated CCT network-tumor system (CNTS) for the evolution of tumor systems. The CNTS activities and behaviors include: uncontrolled cancer cell proliferation, invasion, tumor development (localized disease, regional disease, and metastatic colonized disease), multiple directional dissemination, genome instability, genome diversity, dormancy, new tumor growth, conventional drug pan-resistance, tumor relapse, angiogenesis, and leading to normal tissue and organ biological function failure^1,20–22^.

Briefly, CNTS explains clinical observations and expands the understanding of the nature of cancers: 1) in CNTS, cancer cells do not need to fight against numerous physical ECM and neighbor-cell obstacles that bar metastasis, but can freely migrate in CCTs in the absence of these obstacles; 2) primary-niche cancer cells can directly and efficiently reach far distance destinations via the previously invisible CCT networks and act independent of the circulation systems; 3) disseminated cancer cells can locate or “seed” in many CCT termini and grow into new tumors; 4) CCT membrane superdefences protect cancer cells and tumors inside against conventional cancer drugs and therapies^1,21–22^; 5) CCTs can invade into bone marrows and create new niches difficult for drug therapies (will be reported in another manuscript); 6) CCTs and networks can invade and go through blood-brain barriers and bi-directionally conduct into and outside of the brain (will be reported in another manuscript).

Here, we have comprehensively investigated whether the prevalently employed mouse experimentations of CDX and PDX designed and based on conventional hypotheses for preclinical therapeutic development generate CNTS. We found that there are no CCTs or CCT networks in all these examined CDX and PDX tumors. This study systematically and comprehensively demonstrated the need to terminate conventional animal experimentations and replace with preclinical experimentations engendering CNTS (such as CCT xenograft, CCTX) for the discovery of effective and efficient cancer drugs.

## Results

### Cancer cell line derived xenograft (CDX) models do not generate CCT network-tumor system (CNTS)

Cancer cell line derived xenografts (CDX) in mice are convenient and frequently employed preclinical experiments in the animal models for cancer drug discovery and development^12^. Injections of cancer cells directly into the circulation system are popularly used for the tumor organ colonization^19^; however, this should not be considered a mimic of cancer metastasis. We investigated CDX transplanted tumor tissues of 8 kinds of cancers in breast (MCF-7 breast cancer cells), bladder (HT-1376 bladder cancer cells), colon (HCT116 colon cancer cells), lung (A427 lung cancer cells), pancreas (Bxpc3 pancreas cancer cells), prostate (LnCap prostate cancer cells), ovary (OVCAR3 ovary cancer cells), stomach (NCI-N87 gastric cancer cells). These are within the top 10 most popular and aggressive human cancers assayed via the indicated injection methods (left cardiac ventricle or tail vein). Using histochemistry fluorescence staining assays with anti-CM-01 antibodies and fluorescence microscope, we comprehensively analyzed the CCTs in all the specimens of these 16 CDX transplanted tumors colonized in lung or liver organs (**Fig. 1A**). Consistently, we found that there are no cytocapsular tubes (CCTs) or CCT networks in all these examined 8 kinds of CDX tumor samples (0% of all these samples show CCTs) (**Fig. 1B**). These data show that organ colonized CDX tumors do not generate CCTs, CCT networks, or CNTS in these experimentation models. These experiments demonstrated that cancer drug candidates selected by CDX model in mice, in which CCTs and networks and CNTS are absent, will not have the ability to effectively eliminate CCTs and networks by design, and will meet with conventional cancer drug pan-resistant tumor metastasis (cdp-rtm), which are consistent with the poor outcomes of many cancer drugs selected by these methods.

**Fig. 1.**
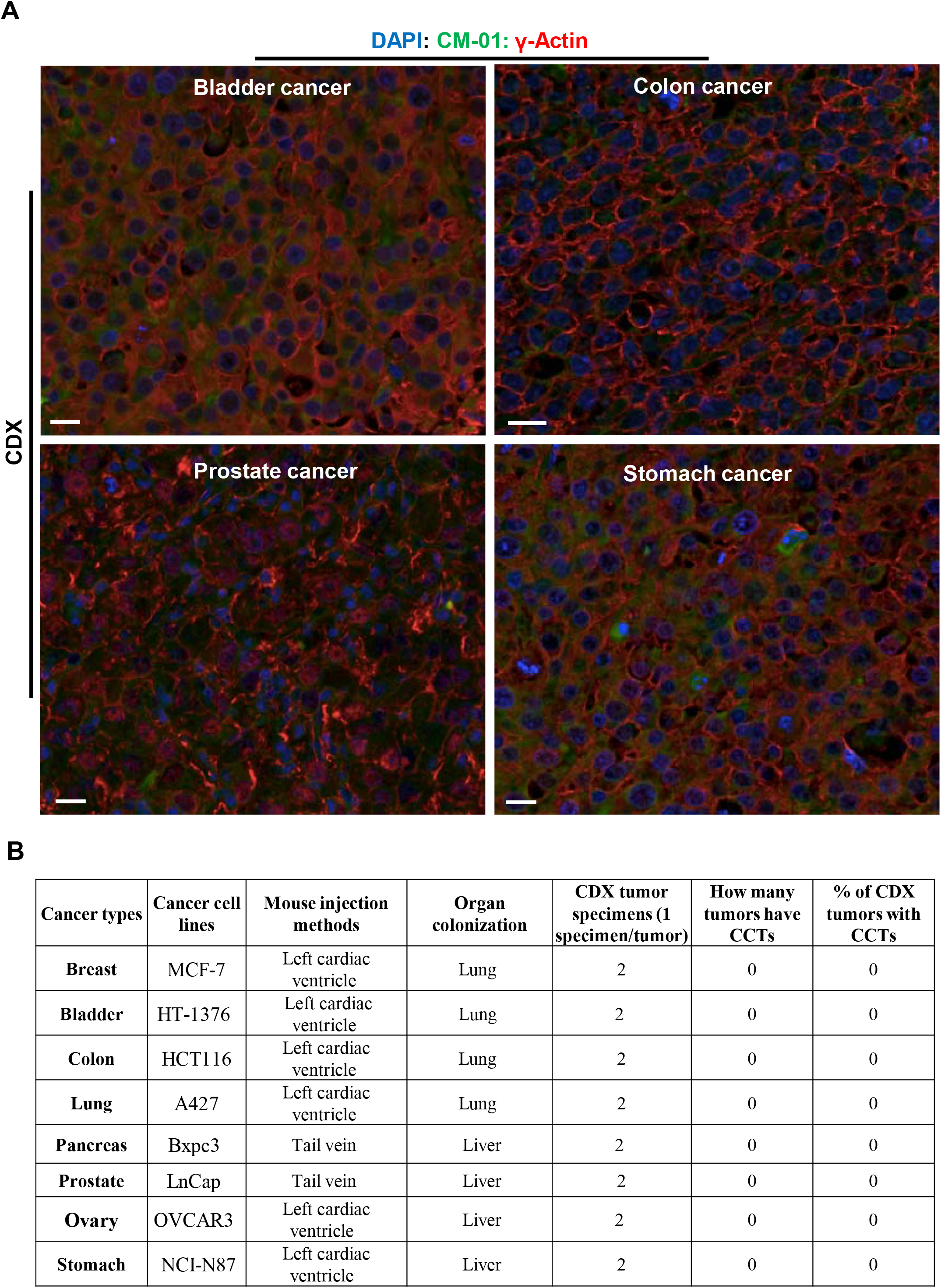
Cancer cell line derived Xenograft (CDX) models do not generate cytocapsular tubes (CCTs), CCT networks or CNTS. **(A)** Representative fluorescence microscope image of CDX transplanted tumor FFPE tissues of cancers in bladder (HT-1376 bladder cancer cell line), colon (HCT116 colon cancer cell line), prostate (LnCap prostate cancer cell line), and stomach (NCI-N87 gastric cancer cell line). There are no CCTs, CCT networks, or CNTS in all these tumor samples. Scale bar, 10μm. **(B) Table of characterization of CCTs in CDX tumors in mice with 8 kinds of cancers with cancer cell lines**.

### Clinical primary malignant tumor cells in patient cancer cell derived xenograft (PDX) models do not engender CNTS

Patient cancer cell line derived xenografts (PDX) in mice are also frequently applied for preclinical experiments in animal models. Primary malignant tumor cells in cancer stage III are propertied with established potent capacities for cancer metastasis in patient bodies *in vivo*^8^. Next, we investigated PDX models (injection methods: left cardiac ventricle or tail vein in mice) with primary malignant tumor cells of 7 kinds of late-stage tumors from patients with cancers: bladder (Stage IIIa), gallbladder (Stage III), Head/neck (Stage IIIc), intestine (Stage IIIa), pancreas (Stage III b), rectum (Stage III), and testis (Stage III). Using histochemistry fluorescence staining assays with anti-CM-01 antibodies and fluorescence microscope, we systematically analyzed the CCTs in all these tumor specimens of the indicated 9 PDX transplanted tumors colonized in lung or liver organs in mice. Consistently, there are no CCTs, CCT networks, or CNTS in the 7 kinds of PDX models with patient primary tumor cells of bladder, gallbladder, Head/neck, intestine, pancreas, rectum, and testis at late cancer stages (none of the samples present CCTs) (**Fig. 2 and Table 1**). These representative and unbiased data evidenced that organ colonized PDX tumors with patient primary tumor cells do not generate CCTs, CCT networks, or CNTS in these animal models. These experimental data demonstrated that cancer drug candidates selected by PDX model (primary tumor cells) in animals, in which CCTs and networks and CNTS are absent, will not have the capacity to efficiently penetrate CCT membranes, overcome the CCT superdefence, kill cancer cells in CCTs, and eliminate CNTS. These data are consistent with the poor outcomes of many cancer drugs developed by these methods.

**Fig. 2.**
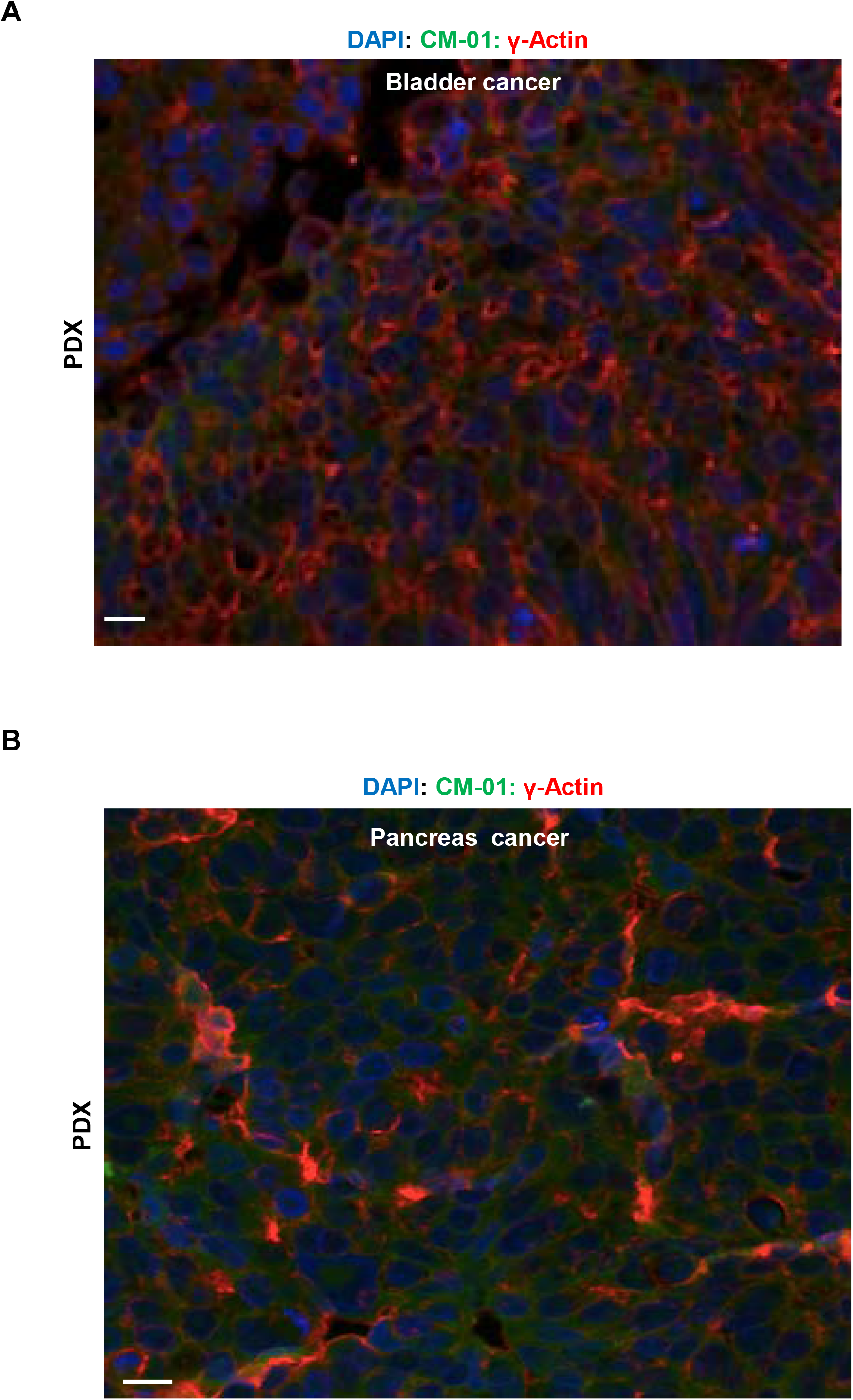
Patient cancer cell derived Xenograft (PDX) models with primary malignant tumor cells via circulation injection and colonized in lung and liver do not generate cytocapsular tubes (CCTs), CCT networks or CNTS. **(A)** Representative fluorescence microscope image of PDX transplanted FFPE tumor tissues with patient (cancer Stage IIIa) bladder primary malignant tumor cells via left cardiac ventricle injection and colonized in lung. There are no CCTs, CCT networks, or CNTS in the indicated tumor samples. Scale bar, 10μm. **(B)** Representative fluorescence microscope image of PDX transplanted FFPE tumor tissues with patient (cancer Stage IIIb) pancreas primary malignant tumor cells tail vein injection and colonized in liver. There are no CCTs, CCT networks, or CNTS in the indicated tumor samples. Scale bar, 10μm.

**Table 1:** Characterization of CCTs and drug resistance in PDX tumors in mice with clinical patient cancer cells from primary tumors and metastatic tumors of 16 kinds of cancers.

| Cancer types (subtypes) | Cancer Stage at the surgery | Primary or metastatic site tumor cells used for PDX | After surgery, cancer patient take how many chemotherapies /show resistance to how many drugs | Drug resistance percentage (%) | If cancer patient die after several chemotherapies | Mouse injection methods | Organ colonization of the PDX transplanted tumors in mouse | Checked PDX transplanted tumor number (1 specimen /tumor) | How many tumors have CCTs | % of CDX tumors with CCTs |
| --- | --- | --- | --- | --- | --- | --- | --- | --- | --- | --- |
| Bladder | IIIa | Primary | 2/2 | 100 | N/A | Left cardiac ventricle | Lung | 2 | 0 | 0 |
| Breast (Triple negative) | IIIc | Metastatic to lung | 3/3 | 100 | N/A | Left cardiac ventricle | Lung | 2 | 0 | 0 |
| Colon | III b | Metastatic to stomach | 2 /2 | 100 | Yes | Tail vein | Liver | 2 | 0 | 0 |
| Gallbladder | III | Primary | 3/3 | 100 | N/A | Left cardiac ventricle | Lung | 1 | 0 | 0 |
| Head/neck | IIIc | Primary | 3/3 | 100 | Yes | Left cardiac ventricle | Liver | 1 | 0 | 0 |
| Intestine | III a | Primary | 2/2 | 100 | N/A | Tail vein | Liver | 2 | 0 | 0 |
| Lung | IV | Metastatic to breast | 4/4 | 100 | Yes | Left cardiac ventricle | Lung | 2 | 0 | 0 |
| Oesophagus | III | Metastatic to lymph | 2/2 | 100 | N/A | Tail vein | Lung | 1 | 0 | 0 |
| Ovary | III | Metastatic to lymph | 2/2 | 100 | N/A | Tail vein | Lung | 2 | 0 | 0 |
| Pancreas | III b | Primary | 3/3 | 100 | Yes | Tail vein | Liver | 1 | 0 | 0 |
| Prostate | III | Metastatic to bladder | 2/2 | 100 | N/A | Left cardiac ventricle | Lung | 2 | 0 | 0 |
| Rectum | III | Primary | 2/2 | 100 | N/A | Left cardiac ventricle | Liver | 1 | 0 | 0 |
| Stomach | III | Metastatic to lung | 2/2 | 100 | N/A | Tail vein | Liver | 1 | 0 | 0 |
| Testis | III | Primary | 2/2 | 100 | N/A | Left cardiac ventricle | Lung | 1 | 0 | 0 |
| Thyroid | III | Metastatic to lymph | 2/2 | 100 | N/A | Left cardiac ventricle | Lung | 1 | 0 | 0 |
| Uterus | IIIc | Metastatic to lymph | 2/2 | 100 | Yes | Left cardiac ventricle | Liver | 1 | 0 | 0 |

### Clinical metastatic tumor cells in PDX models do not engender CNTS

Clinical metastatic cancer cells generate a lot of CCTs and networks and CNTS in cancer patient tissues and organs. Metastatic cancer cells are propertied with potent capacities of engendering CCTs and networks. Thus, we investigated PDX models (injection via left cardiac ventricle or tail vein) with metastatic secondary tumor cells of multiple late stage tumors from patients with cancers: breast (tripe negative, lack of expression of estrogen receptor (ER), progesterone receptor (PR), and human epidermal growth factor receptor-2 (HER2); Stage IIIc; in patient lung), colon (Stage III b; in patient stomach), lung (Stage IV; in patient breast), oesophagus (Stage III, in patient lymph node), ovary (Stage III; in patient lymph node), prostate (Stage III; in patient bladder), stomach (or gastric, Stage III, in patient lung), thyroid (Stage III, in patient lymph node), and uterus (Stage IIIc; in patient lymph node). Then, we comprehensively analyzed the CCTs in all these tumor specimens of the indicated 13 PDX transplanted tumors colonized in mice lung or liver (**Fig. 3 and Table 1**). Consistently, we found that there are no CCTs, CCT networks, or CNTS in all 9 kinds of PDX models with patient metastatic secondary tumor cells of breast, colon, lung, oesophagus, ovary, prostate, stomach, thyroid, and uterus at late cancer stages (none of the samples display CCTs). These comprehensive data display that organ colonized PDX tumors of patient metastatic tumor cells do not generate CCTs, CCT networks, or CNTS in these animal models. The aforementioned experimental results demonstrated that cancer drug candidates selected by PDX model (with metastatic tumor cells) in animals, in which CCTs, networks and CNTS are absent, will have poor or no capacity to efficiently penetrate CCT membranes, overcome the CCT superdefence, kill cancer cells in CCTs, and eliminate CNTS by design. These observations are consistent with the poor outcomes of many cancer drugs developed by these models.

**Fig. 3.**
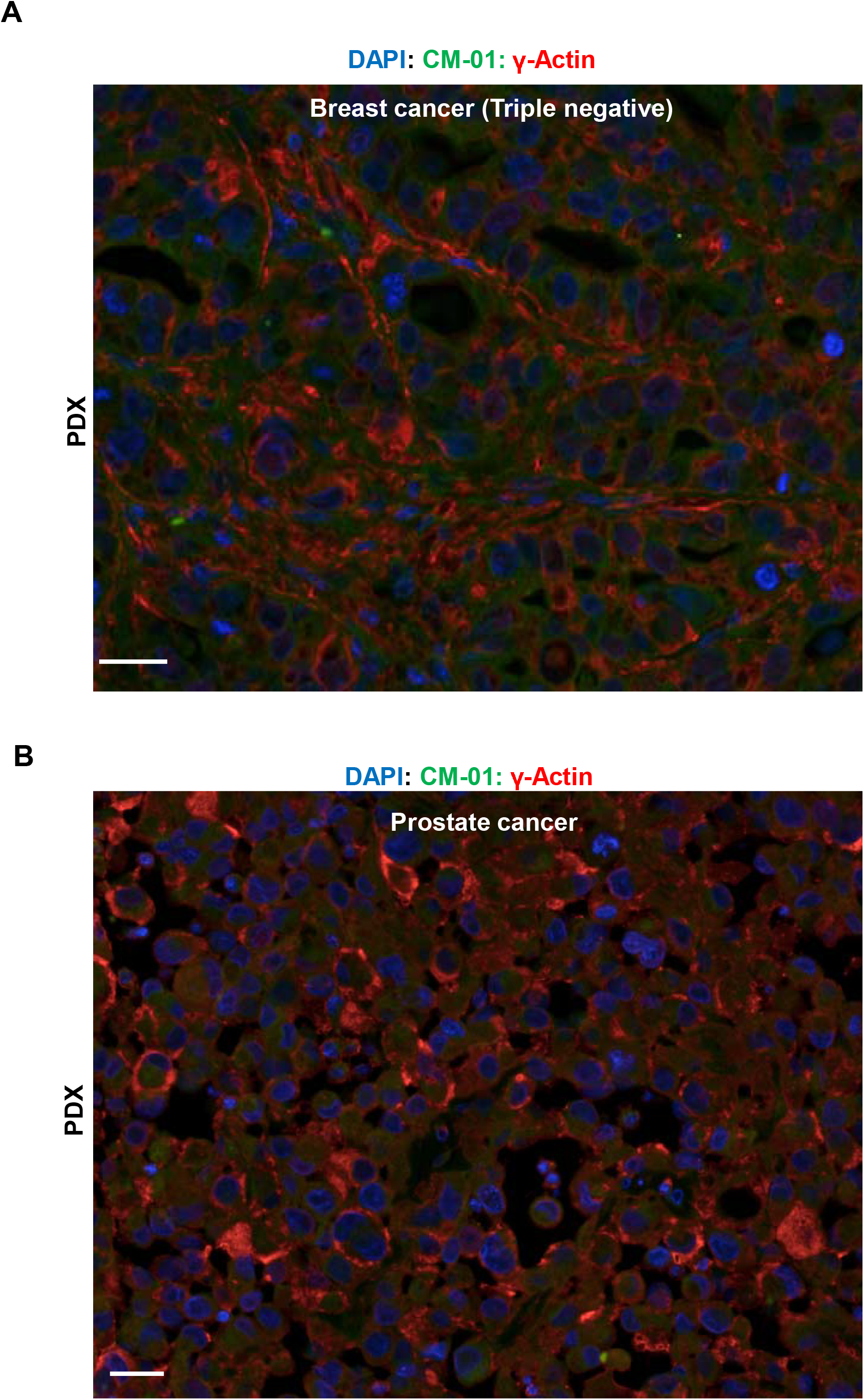
Patient cancer cell derived Xenograft (PDX) models with metastatic secondary malignant tumor cells via circulation injection and colonized in lung do not generate cytocapsular tubes (CCTs), CCT networks or CNTS. **(A)** Representative fluorescence microscope image of PDX transplanted FFPE tumor tissues with patient (cancer Stage IIIc), triple negative (ER-/PR-/HER2-) breast metastatic secondary malignant tumor cells via left cardiac ventricle injection and colonized in lung. There are no CCTs, CCT networks, or CNTS in the indicated tumor samples. Scale bar, 10μm. **(B)** Representative fluorescence microscope image of PDX transplanted FFPE tumor tissues with patient (cancer Stage III) prostate metastatic secondary malignant tumor cells via left cardiac ventricle injection and colonized in lung. There are no CCTs, CCT networks, or CNTS in the indicated tumor samples. Scale bar, 10μm.

In summary, the above comprehensive experimentation data evidenced that conventional CDX and PDX animal models do not generate CCT networks or CNTS. These data demonstrated that cancer drugs developed by these models will have poor or no ability to effectively and efficiently penetrate CCT membranes, ruin CCT superdefences, kill cancer cells in CCTs, and eradiate CNTS in clinical cancer pharmacotherapy.

### Conventional cancer drugs developed by CNTS absent models/methods have low therapy effectiveness and efficiency in clinical cancer pharmacotherapy

CNTS induced conventional cancer drug pan-resistant tumor metastasis (cdp-rtm) is a major source of cancer death. Next, we investigated the drug resistance of the indicated cancer patients by examining the resistance percentages to the chemotherapy drugs. We found that all (100%) cancer patients in this study show drug resistance to all of the applied conventional cancer drugs with exhibiting 100% drug resistance. Furthermore, the patients with cancers in colon, head/neck, lung, pancreas, and uterus in this study died within 1 year after several chemotherapies with cancer drugs developed by conventional tests and animal models (**Table 1**). The above results, combined with the data that approximately 10million cancer deaths worldwide every year with high percentages of them took chemotherapies by variable numbers of cancer drugs^2^, undoubtedly evidenced that the conventional cancer drugs developed by traditional animal methods without CCTs, CCT networks or CNTS will not have the ability to effectively or efficiently cure solid cancers with superlarge CNTS.

In summary, the aforementioned data demonstrated that CNTS is an integrated physical target for pharmacotherapy, and that preclinical experimentations generating CNTS (such as CCT xenograft, CCTX) (**Fig. 4**) need to be employed for the development of highly effective and efficient cancer drugs aimed for the cure of solid cancers.

**Fig. 4.**
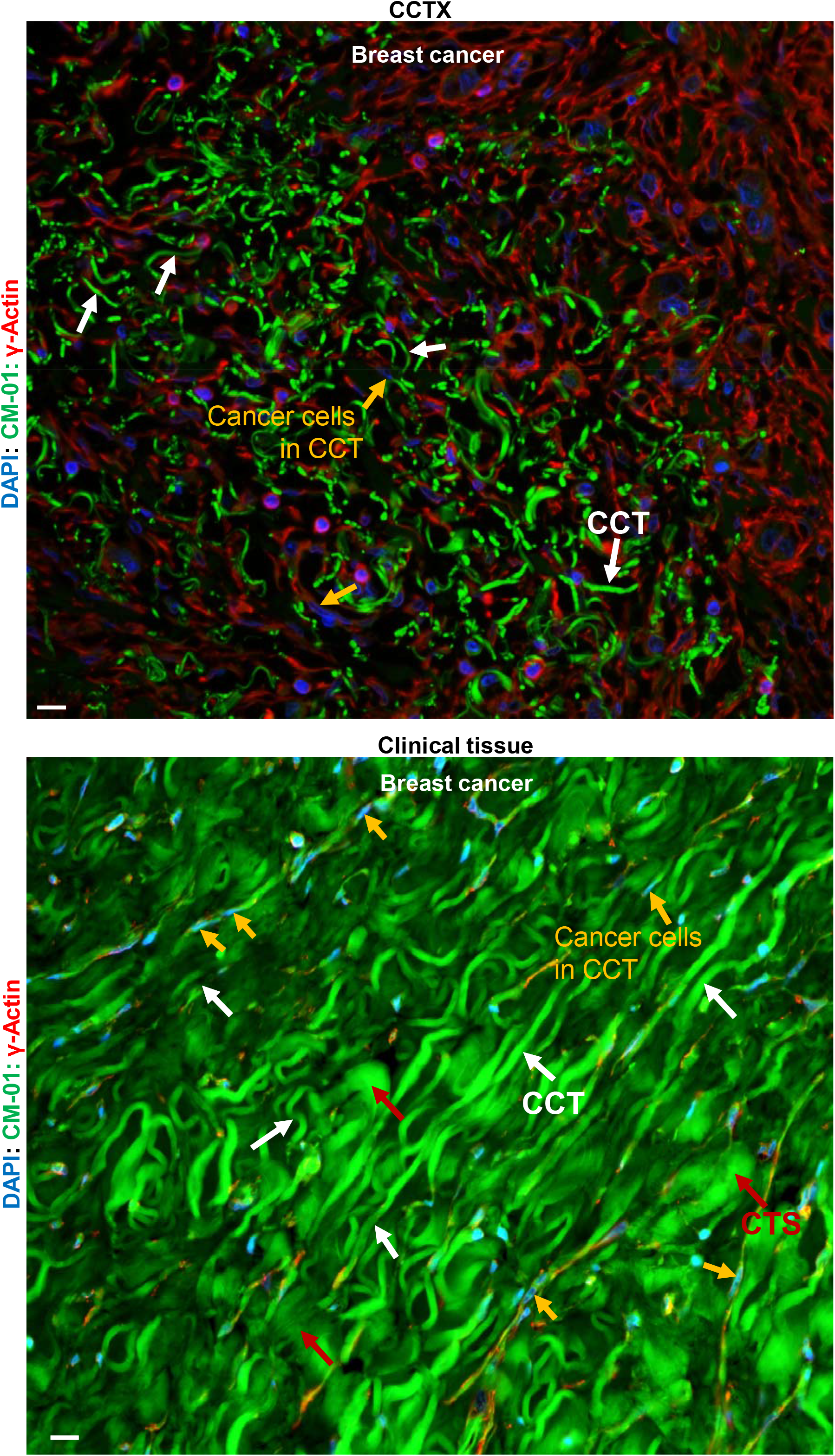
Cytocapsular tube (CCT) Xenograft (CCTX) model and clinical cancer tissues generate a large quantity of CCTs, CCT networks and CNTS. **(A)** Representative fluorescence microscope image of CCTX transplanted FFPE tumor tissues with patient breast tumor cells of CH-5^high^/CH-6^high^ subpopulation breast cancer cells via subcutaneous injection. There are a lot of CCTs (many are sectioned as curved fragments) in these CCTX breast tumor samples. Many CCTs interconnect and form CCT networks. Breast cancer cells migrate in CCTs. The CCTs, CCT networks, breast cancer cells in migration in CCTs, and CCT clusters/masses in the transplanted tumor block (even no secondary metastatic tumor formation in mice in this study) have successfully formed and established small scale and complex CCT network-tumor system (CNTS), which functionally reconstructed the CNTS in transplanted tumors in animal models. Cytocapsular tube (CCT, white arrows), and cancer cells in migration in CCTs (orange arrow) are shown. Scale bar, 10μm. **(B)** Representative fluorescence microscope image of FFPE clinical breast tumor tissues (cancer Stage III). There are countless straight or curved CCTs, which pile up and form CCT masses in these tumor samples. Large quantities of CCTs interconnect and form superlarge CCT networks. Breast cancer cells migrate in CCTs and CCT networks. The CCTs and CCT networks interconnect most/all tumors, and form superlarge and integrated CCT network-tumor system (CNTS) in the cancer patients. Cytocapsular tube (CCT, white arrows), CCT strand (CTS, red arrows), and cancer cells in migration in CCTs (orange arrow) are shown. Scale bar, 10μm.

## Discussion

The development of effective cancer drugs is essential for the precise and efficient cancer pharmacotherapy^2,5–6^. CCT network-tumor system (CNTS) is an integrated superlarge structure in which cancer evolution, metastasis and cdp-rtm occur^1,21–22^. Here, we investigated CNTS in conventionally employed cancer drug discovery platforms of 8 kinds of CDX models and 16 kinds of PDX models and discovered that CNTS is consistently absent in these 24 kinds of animal models. This comprehensive study demonstrated that conventional animal experimentations without CCTs, CCT networks or CNTS, need to be terminated and replaced by appropriate models that generate CNTS for the discovery and development of highly effective cancer drugs for highly precise and efficient tumor pharmacotherapy.

### The CCT network-tumor system (CNTS) is an integrated organization composed of CCT networks and all tumors in patients

Previous hypotheses thought that: 1) primary and metastatic secondary tumors (cancer blocks) in cancer patients are physically separated, biological-behavior-independent, and locally distributed in different sites without physical interconnection between them; 2) cancerous cells must overcome numerous physical ECM and neighbor-cell obstacles barring cell migration and must borrow vasculature pathways to reach far distance destinations *in vivo*; 3) aggressive surgery of visible tumor blocks clear the cancers with no need for follow-up pharmacotherapy. All these thoughts are controversial to clinical observations and are incorrect. Indeed, in cancer patients, 1) all primary and metastatic secondary tumors are interconnected by CCT networks and form an integrated superlarge physical structure of CNTS composed of all tumors and CCT networks; cancer cells can aggregate into and disassemble from tumor blocks via CCT networks; 2) cancer cells in CCT lumens do not contact physical ECM and neighbor cell obstacles shielded by CCT membranes outside, but freely migrate in membrane-enclosed CCT networks and conveniently reach far destinations across tissues and organs independent of the vasculatures; 3) even after aggressive surgery of visible tumor blocks, there are still large quantities of CCTs, networks, cancer cells in CCT lumens, and invisible small tumors (<0.5mm in diameter/width) remain; and these spared cancer cells are the resources for tumor relapse; 4) CCTs connected to tumor blocks can degrade and temporarily create isolated tumor islands, in which the enlarged cytocapsular tube terminals enclose and form large cytocapsulae wrapping tumors; cytocapsulae can also degrade and generate naked tumors, depending on the local microenvironment situations; these naked tumors can regenerate CCTs and networks and re-interconnect with each other tumors in different sites; so, the CNTS is a dynamic superlarge organization; 5) One type of tissue originated cancer cells can create a kind of CNTS, and different types of tissue originated cancer cells can generate different kinds of CNTSs (CCT protein types, species and abundances are different); different kinds of CNTSs can combine together by the interconnections of different kinds of CCTs and networks^1,21–24^. After radio-, chemo-, immune-, physical-therapies, there are many CCT networks and cancer cells in CCTs remain, which are the resources for cancer relapse. Therefore, the whole CNTS, not only the detectable tumors (by eye, CT-scan or MRI), but also all invisible CCTs, CCT networks, cancer cells in CCTs, tiny tumors, sporadically scattered cancer cell masses that may connect CNTS via CCTs, should be considered as an integrated physical system during cancer therapy and treatment in order to achieve highly effective outcomes.

Therefore, the conventional cytostatic therapies that only interrupt colonization pathways (not the physical pathways in the metastatic cascade hypothesis which is controversial to clinical observations) rather than kill proliferating tumor cells, or the cytotoxic therapies that only kill parts of proliferating tumor cells and shrink tumors but do not eliminate CCTs and networks, will not effectively destroy or eliminate the whole CNTS structure. Thus, the physical and integrated CNTS should be considered as one superlarge and massive target for the effective and efficient solid cancer therapy.

### New methods that generate CNTS (such as CCTX) should be employed for the discovery and development of highly effective and efficient cancer drugs

It is clear that all conventional transplantations of human cancer cells into other species to develop tumors (or mosaic tumors), no matter what kinds of injection methods and colonized tumor sites, do not generate CNTS for efficient cancer drug candidate test and selection. The CCTX models generate a lot of CCTs, CCT networks and cancer cells migration in CCTs, and engender CNTS platform in transplanted tumors in mice^1^. Therefore, CCTX models provide the appropriate platforms for the discovery of highly effective and efficient cancer drug candidates. They will allow efficient clinical trials for developing precise and effectual cancer drugs, and yield high quality outcomes of cancer pharmacotherapy. Assays that contain cancer drug panresistant tumor metastasis (dcp-rtm) physical superdefence freeway systems, cytocapsular tubes, and networks will be needed for the discovery and development of highly effective and efficient cancer drugs.

Animal experimentations for preclinical therapeutic development that do not generate CCTs and networks should be avoided to prevent huge cancer cost, eliminate unjustified animal experiments, prevent synthesis of low efficient drugs, and save approximately 10million cancer patients’ lives worldwide every year. It is unknow why these transplanted tumors do not engender CCTs, CCT networks or CNTS. It is also unknown what elements and factors are necessary and sufficient for the stimulation of CCT generation. It needs to perform more research to elucidate whether CNTSs are present in the genetically engineered mouse (GEM) with spontaneous tumor metastasis or the orthotopic transplantation of the mouse tumors in the tissues of origin.

### CNTS is the druggable physical target for the cure of all kinds of solid cancers

The integrated superlarge CNTS structure makes it a physical target for cancer therapy. There are multiple advantages for targeting the physical and massive CNTS: 1) the superlarge CCT membranes contain a variety of proteins with essential CCT functions, and can be used as molecular targets for therapy; 2) the integration of the protection shield layer of CCT membranes can be targeted; the damage and elimination of CCT fragments will directly affect the cancer cells inside or make these cancer cells more accessible to cancer drugs for inhibition; 3) the block, damage, or ruin of CCTs and CCT networks will eliminate cancer cell dissemination freeways and therefore inhibit cancer metastasis; 4) the inhibition of CCT initiation, elongation, interaction and fusion, cancer cell migration inside CCT lumens, CCT or network maintenance will effectively eliminate cancer metastasis pathways, and therefore efficiently inhibit tumor metastasis and conventional cdp-rtm; 5) cancer cells in CCTs or in tumors in enlarged cytocapsulae can be targeted by disrupting CCT membranes to allow cancer drugs penetrate into the lumens, and therefore access and kill cancer cells.

It is necessary to pay attention to the new challenges on CNTS elimination: 1) physical drug barrier to cancer drugs that does not penetrate or eliminate CCTs; 2) damage-resilience of CCT networks; 3) separation of affected CCTs; 4) density heterogenicity of CCT densities and networks; 5) potent capacities of cancer cells to (re)generate new CCTs; 6) heterogenicity of CCT membrane protein types, species and abundance; 7) acquired drug resistance of CCT membranes; 8) the potent plasticity and flexibility of CCTs, CCT networks and CNTS.

In summary, this study investigated the CNTS in respect to conventional cancer drug developmental animal tumor platforms. It demonstrated that tumor experimentations with CNTS should be employed in order to develop highly effective cancer drugs for the precise and efficient solid cancer pharmacotherapy at the personal and broad-spectrum levels.

## Methods and Materials

### Tissue slides, antibodies, and reagents

Formalin-fixed paraffin-embedded (FFPE) CDX and PDX tissue slides were ordered from Charles River laboratories, US Biomax, or local hospitals. Monoclonal mouse anti-γ-actin antibodies were ordered from Abcam (ab123034). Polyclonal and monoclonal rabbit anti-CM-01 (code, not real protein name) antibodies were self-developed. Secondary antibodies of antimouse and anti-rabbit antibodies were ordered from Thermo Fisher Scientific. DAPI was ordered from VWR. Polyclonal and monoclonal rabbit anti-CH-5 and anti-CH-6 (codes, not real protein names) antibodies were self-developed. Secondary antibodies of anti-mouse and anti-rabbit antibodies were ordered from Thermo Fisher Scientific. DAPI was ordered from VWR. Matrigel matrix was ordered from Sigma or Corning. *NOD-SCID* (strain name: *NOD.CB17-Prkdcscid/J*) were ordered from The Jackson Laboratory or Charles River Laboratories. All mouse work were reviewed and permitted by Cytocapsula Research Institute Animal Committee and performed in Charles River Laboratories.

### Immunohistochemistry (IHC) fluorescence staining assay

The sectioned FFPE clinical cancer tissue specimens were subjected to double immunohistochemistry staining with anti–CM-01 (1:200) and anti–γ-actin (1:200) primary antibodies and DAPI (1:1,000) for 4h at 4 °C, followed by incubation with appropriate secondary antibodies for 1h at 4 °C in a dark room. Fluorescence images were taken with a Nikon 80i upright microscope with a 20× or 40× lens. All images were obtained using MetaMorph image acquisition software and were analyzed with ImageJ software.

### Cell cytometry

Patient breast cancer cells were extracted from the fresh clinical cancer tissues, and cultured with culturing situations mimicking *in vivo* environments: 37°C, 5% CO_2_, humidity and a series of other conditions. The CH-5^high^/CH-6^high^ patient cancer cell subpopulations were isolated by fluorescence-activated cell sorting (FACS) assay with self-developed FITC-conjugated anti-CH-5 and PE-conjugated anti-CH-6 antibodies.

### Tumor xenograft formation assay

In the tumor xenografted assay, the indicated patient cancer cells, and CH-5^high^/CH-6^high^ subpopulation breast cancer cells (cell numbers: 0.5million) were mixed with 100μl Matrigel/cell culture media mixture (Matrigel: cell culture media = 1:2) (Sigma or Corning). Indicated cancer cells/Matrigel/ cell culture media mixtures were injected into *NOD/SCID* female for breast cancer xenograft (the Jackson Laboratory) by subcutaneous injection. After the tumor formation (about 75 mm^3^ in volume, 5 mice/group), mice were sacrificed and tumors were excised. Tumor tissue samples were used for immunohistostaining and CCT analyses. The mouse experiments were performed according to the policies of Cytocapsula Research Institute Animal Committee and Charles River Laboratories.

### Statistical analysis

Quantitative data were statistically analyzed (mean ± SD, *t*-test, two-tailed). Statistical significance was determined by *t*-test. Significance was expressed as: *:*p*< 0.1; **:*p*< 0.05; ***: *p*< 0.01, or with the *p*-value. *P*<0.05 was considered significant.

Supported by a grant (to Dr. Yi) from Cytocapsula Research Institute Fund for Cytocapsular Tube Conducted Cancer Metastasis Research, and a grant (to Dr. Yi) from Chalst Inc Fund for Cytocapsular Tube Cancer Metastasis Research.

We greatly acknowledge Dr. Ed Harlow of Harvard Medical School (USA) and Cancer Institute of University of Cambridge (UK), and Dr. Nahum Sonenberg of McGill University of Canada, Dr Yubo Yang, Dr. Qiping Hou, and Mr. James Ranieri for their help and meaningful discussion in the study.

